# Effects of sequence splitting and negative-set selection on antimicrobial peptide prediction

**DOI:** 10.64898/2026.11.20.768089

**Authors:** Chen Chen, Hongbo Zhou, Qifu Li, Zishang Gao, Yanshan Tian

## Abstract

Sequence partitioning and negative-set selection determine the prediction problem posed by an antimicrobial peptide (AMP) benchmark. We quantified their effects using a DRAMP 5.0 audit, controlled comparisons with frozen ESM-2 classifiers and additional benchmark configurations. On DRAMP, random splitting increased Matthews correlation coefficient (MCC) from 0.718 under cluster-disjoint splitting to 0.932. This difference of 0.214 exceeded the gains from the architectural additions tested, and 75.6% of randomly assigned test sequences shared a homology cluster with training sequences. Across three additional dataset configurations, random-minus-cluster MCC differences ranged from −0.04634 to +0.05682, showing that the magnitude and direction of the split effect depended on benchmark composition. With the fitted classifiers and APD6 positives held fixed, replacing DRAMP test negatives with external hard negatives reduced the area under the receiver operating characteristic curve from 0.9833 to 0.9018 (paired difference 0.08149, bootstrap 95% CI 0.07196 to 0.09181). These comparisons quantify substantial sensitivity to evaluation design and show why split effects require assessment within each benchmark. AMPBenchAudit provides the audited records, split assignments and evaluation code needed to examine this sensitivity and interpret model comparisons under explicit test conditions.

**Author summary:** Antimicrobial peptides are short chains of amino acids that can inhibit microbes. Computer models help researchers select peptides for laboratory testing, and test scores are often used to compare these models. We examined how those scores change when researchers change the examples used for training and evaluation.

Using an audited peptide database, we found that a model scored substantially higher after a random division of the data. About three quarters of its test peptides then had a close relative in the training set. This score difference exceeded the improvements from the architectural additions we tested. Repeating the comparison in three other test setups produced smaller differences, sometimes in the opposite direction. We also kept the model and known antimicrobial peptides fixed while changing the comparison peptides labelled as inactive. The model’s ability to distinguish the groups declined.

These experiments show how benchmark construction influences the evidence used to judge peptide predictors. Our openly available data and code allow researchers to inspect sequence overlap and repeat the comparisons with clearly documented test examples.

## Introduction

Antimicrobial peptide prediction is usually posed as a binary sequence-classification task, with activity labels taken from curated databases and performance reported as accuracy, area under the receiver operating characteristic curve (AUROC) or MCC on a held-out set. The field has accumulated many such reports [1,2,3,4,5], and the models have grown more elaborate, from composition-based classifiers to protein language models and graph neural networks [6,7,8,9]. Recent work has called for standard evaluation frameworks and shared benchmarks for AMP activity data [10,11]. Comparing the numbers across papers is nevertheless unreliable. Studies use different source releases, different rules for resolving multiple activity measurements, different negative sets, and different ways of dividing sequences into training and test data. These concerns are not specific to peptides: audits of machine-learning benchmarks have documented leakage and split sensitivity in protein machine learning and other domains [12,13,14,15].

Curated AMP resources contain short, closely related sequences. A random split can place near-duplicates in training and test sets. Several lines of work have argued that evaluation on homology-aware partitions gives a more realistic estimate of generalisation [12,14,16], and the size of the resulting score difference can be measured by comparing split schemes within a dataset. Negative-set provenance also matters in AMP evaluation [10]. When positive and negative labels come from different databases, a classifier may learn features associated with the source.

The data themselves add a third complication. DRAMP 5.0 provides activity annotations and FASTS exports that combine peptide sequences with SMILES [17]. In the records we audited, minimum inhibitory concentration (MIC) values required parsing from free-text fields, and some structure annotations disagreed with the recorded sequence. Curation conventions differ across the major peptide resources [18,19,20,21], and MIC determination itself is protocol dependent [22]. Without a record-level audit, a modelling pipeline can inherit endpoint mixing, duplicated measurements and mismatched structure annotations before any split is made. Variation between repeat measurements gives an indication of how consistently the regression target was measured.

We asked how much AMP performance estimates change with sequence partitioning and negative-set selection, and whether the split effect persists across benchmark configurations. AMPBenchAudit combines a record-level DRAMP 5.0 audit with comparisons that hold the representation and training protocol fixed while changing the split scheme. A complementary experiment holds the fitted classifiers and APD6 positives fixed while changing the source of test negatives. Together, these experiments quantify the sensitivity of reported performance to two common evaluation choices. Comparisons across additional datasets establish how that sensitivity varies with benchmark composition. Public-tool evaluations, architectural ablations and external MIC analyses provide context for the size of these effects and the transfer of predictions across data sources. The released records, split assignments and code support reproduction of each comparison.

## Results

### Record audit and construction of the prediction cohorts

DRAMP 5.0 contains 32,296 records, of which 22,344 carry a SMILES string (Fig 1, Table 1, S1 Fig). Graph isomorphism between the SMILES-derived structure and the FASTA sequence was observed in 35.4% of general records and 6.6% of stability records. Patent-derived records dominated the pooled estimate of 84.9%, so we analysed the curated subsets separately. We parsed MIC values from the free-text organism field, and 0.4% of organism-name entries remained unresolved. Four rules for aggregating repeat measurements agreed at *ρ≥* 0.978. The Spearman Brown corrected consistency was approximately 0.82 among records with repeat measurements.

**Table 1.** Data audit and cohort construction.

| **Quantity** | **Value** |
|---|---|
| DRAMP 5.0 records | 32,296 |
| Records carrying a SMILES string | 22,344 |
| SMILES–sequence graph isomorphism (general / stability / pooled) | 35.4% / 6.6% / 84.9% |
| Organism-name entries unresolved | 0.4% |
| Agreement across MIC aggregation rules | *ρ*≥0.978 |
| Spearman–Brown label consistency | ≈0.82 |
| Negative set | 13,204 rows (5,886 unique) |
| Task-1 split | 26,408 sequences (test 2,549) |
| Task-3 split | 2,378 sequences (test 238) |

**Fig 1.**
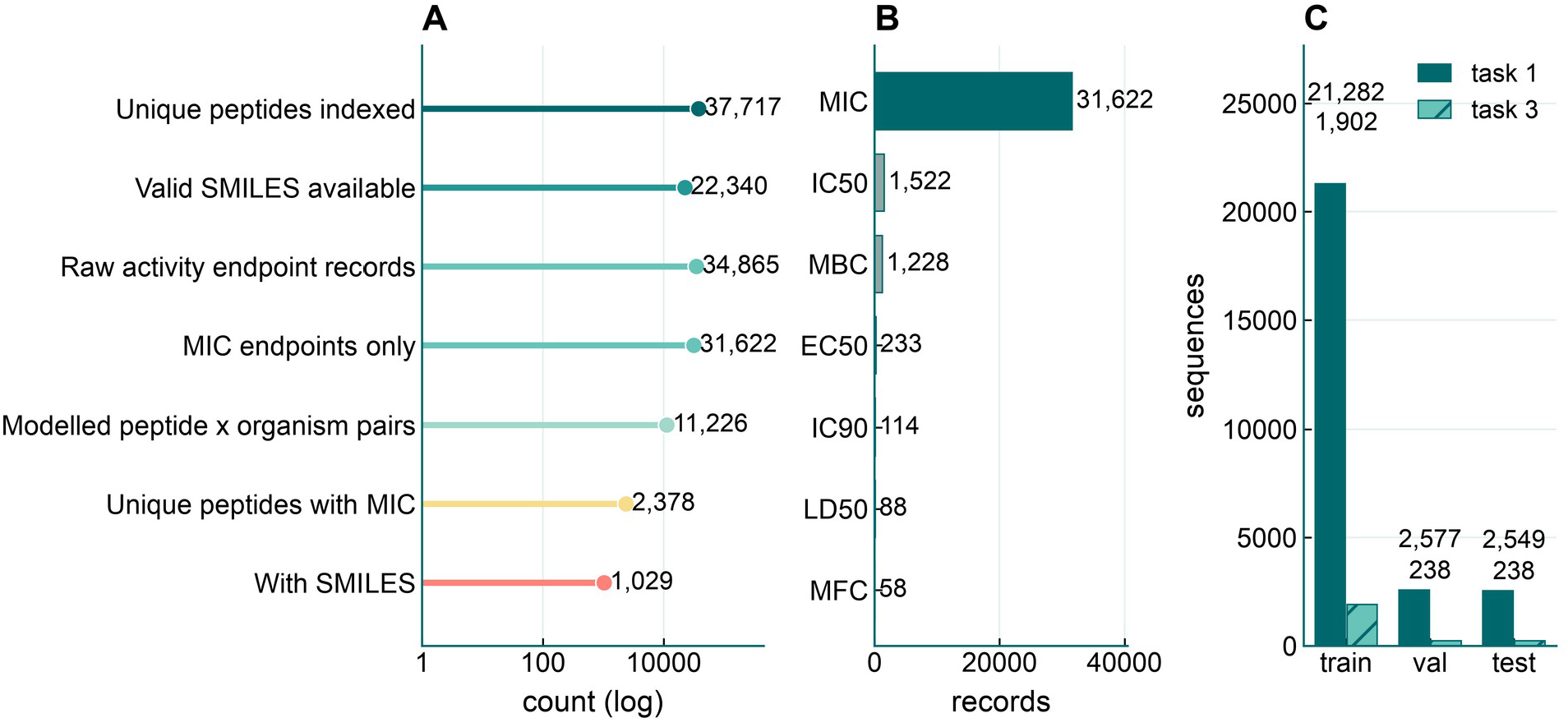
Data audit and cohort construction. (A) Counts from the sequence index and activity audit, shown on a logarithmic scale. The index contained 37,717 peptide entries, of which 22,340 had a valid SMILES representation. Of 34,865 activity records, 31,622 reported a minimum inhibitory concentration (MIC). The resulting regression cohort contained 11,226 peptide–organism pairs across 2,378 peptide entries, of which 1,029 had SMILES. (B) Counts of activity records by endpoint. Only MIC measurements contributed to the regression target. The other recorded endpoints were IC50, MBC, EC50, IC90, LD50 and MFC. (C) Sequence counts in the training, validation and test sets for AMP classification (task 1, solid bars) and pMIC regression (task 3, hatched bars). For each task, we clustered sequences by single linkage at a sequence identity threshold of 0.80 and assigned whole clusters to one partition each. The upper labels show task 1 counts (21,282, 2,577 and 2,549), and the lower labels show task 3 counts (1,902, 238 and 238).

Values are taken from the frozen audit tables and split files; the two split files are not interchangeable. The pooled isomorphism figure is dominated by patent-derived records and was not used for modelling decisions.

The audit yielded 13,204 negative-set rows (5,886 unique sequences), a task-1 partition of 26,408 sequences with 2,549 held out for test, and a separate task-3 partition of 2,378 sequences with 238 held out for test. Every figure and number in this paper is drawn from that snapshot; no result was recomputed after the freeze.

### Fixed-threshold comparison with public AMP predictors

With a fixed decision threshold of 0.5 and five seeds, the frozen ESM-2 representation plus a shallow head gave MCC 0.7204 ± 0.0204 and mean per-seed AUROC 0.9404 (Fig 2, Table 2). Running four public predictors [23,24,25,26] on the same partition gave 0.699 for AMPlify, 0.613 for Macrel, 0.557 for AMP Scanner v2 and 0.483 for AMP-BERT. On the paired subset used for inference (n = 2,225), MCC was 0.71945 against 0.69859 for AMPlify, a mean per-seed difference of +0.0209. The paired bootstrap of the five-model mean prediction score gave a 95% CI of 0.0082 to 0.0752 for the corresponding ensemble MCC difference. The per-seed mean difference was below the prespecified margin of 0.03, and the ensemble interval spans that margin. On the same subset, AUROC based on the ESM-2 mean prediction score differed by +0.0270 (DeLong 95% CI 0.0163 to 0.0377, *p* = 7.22 *×* 10^-7^). Table 3 reports the number of matched test sequences and MCC for each predictor pair. S2 Fig gives per-seed and threshold-sensitivity results. Macrel’s output omitted the N-terminal methionine in 423 of 2,196 records. Matching by exact sequence yielded 1,790 rows (MCC = 0.613), and matching by record ID yielded 2,196 rows (MCC = 0.635).

**Table 2.** Fixed-threshold benchmark against four public predictors.

| **Predictor** | **MCC at 0.5** | **n** |
|---|---|---|
| Frozen ESM-2 representation with a shallow head | 0.7204 ± 0.0204 | 2,549 |
| AMPlify | 0.699 | 2,549 (paired 2,225) |
| Macrel | 0.613 / 0.635 | 1,790 / 2,196 |
| AMP Scanner v2 | 0.557 | 2,221 |
| AMP-BERT | 0.483 | 2,549 |

Predictors were evaluated on the DRAMP test partition using the rows available for each tool. The five-seed spread is for the ESM-2 classifier. For Macrel, the descriptive results use sequence-key and ID-key alignments, respectively.

**Table 3.** Matched pairwise comparisons against public predictors. The denominator for coverage is the full test set of 2,549 rows. Each row compares the two predictors on the same sequences.

| **Public predictor** | **Matched n (coverage)** | **ESM-2 MCC** | **Public predictor MCC** | **ΔMCC** |
|---|---|---|---|---|
| AMPlify | 2,225 (87.3%) | 0.71945 | 0.69859 | +0.02086 |
| Macrel (ID key) | 2,196 (86.2%) | 0.71756 | 0.63527 | +0.08229 |
| AMP Scanner v2 | 2,221 (87.1%) | 0.73131 | 0.55685 | +0.17446 |
| AMP-BERT | 2,549 (100.0%) | 0.72039 | 0.48309 | +0.23731 |

**Fig 2.**
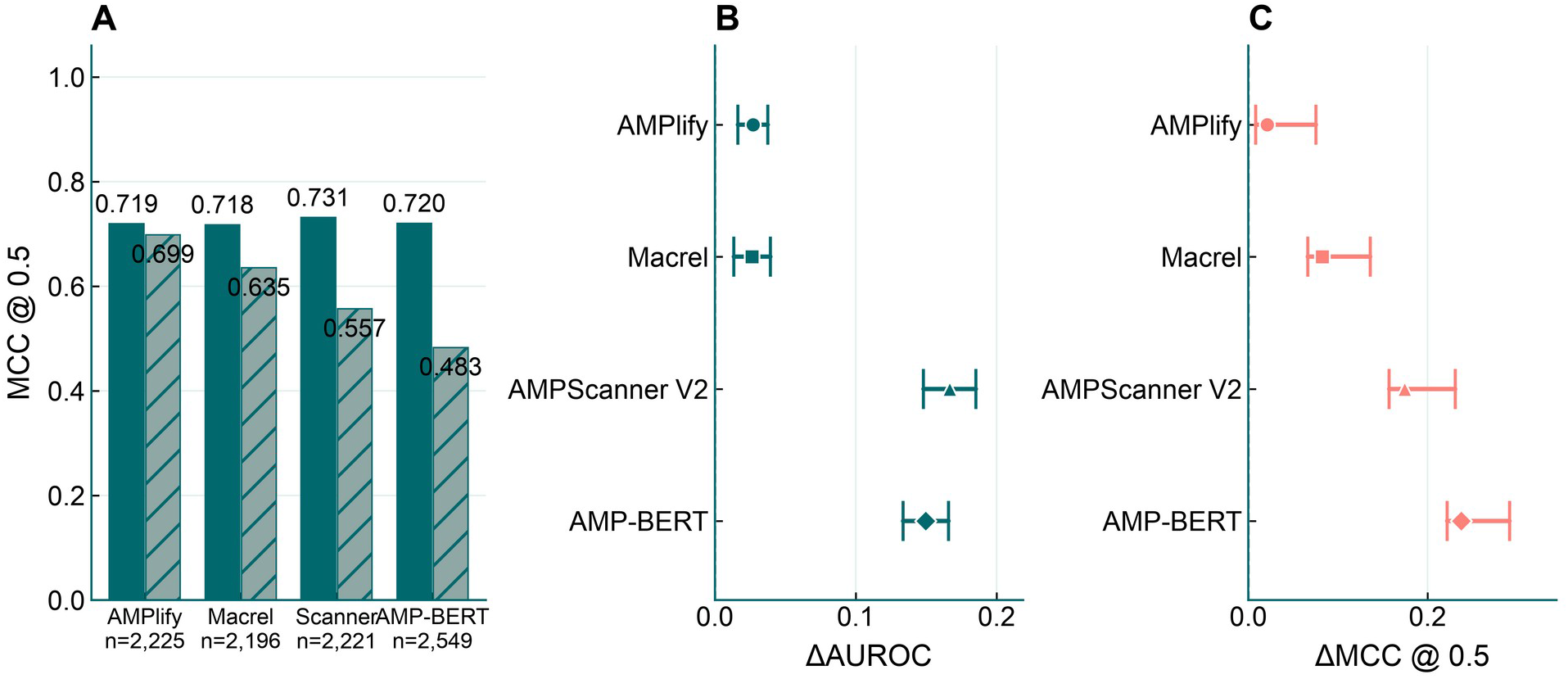
Fixed-threshold benchmark against public tools. (A) Matthews correlation coefficient (MCC) at threshold 0.5 on the matched test subset for each public predictor. Solid bars show the mean MCC across five ESM-2 runs. Hatched bars show public predictor scores. Sample sizes are shown below the bars. (B) Paired AUROC differences between the five-run mean ESM-2 prediction and each public predictor, with 95% DeLong confidence intervals. (C) Differences in MCC at threshold 0.5 for the same comparisons. Points show gaps based on mean per-run MCC, and intervals are paired bootstrap 95% confidence intervals based on mean prediction probabilities.

### Classification scores under two DRAMP split schemes

We used global sequence alignment [27] to compute sequence identity, applied single-linkage clustering at a threshold of 0.80 and assigned whole clusters to the training, validation and test partitions (Fig 3). Under random splitting, 75.6% of test sequences had a homologue in training. MCC increased from 0.718 with the cluster-disjoint split to 0.932 with the random split, a difference of 0.214 on this cohort.

**Fig 3.**
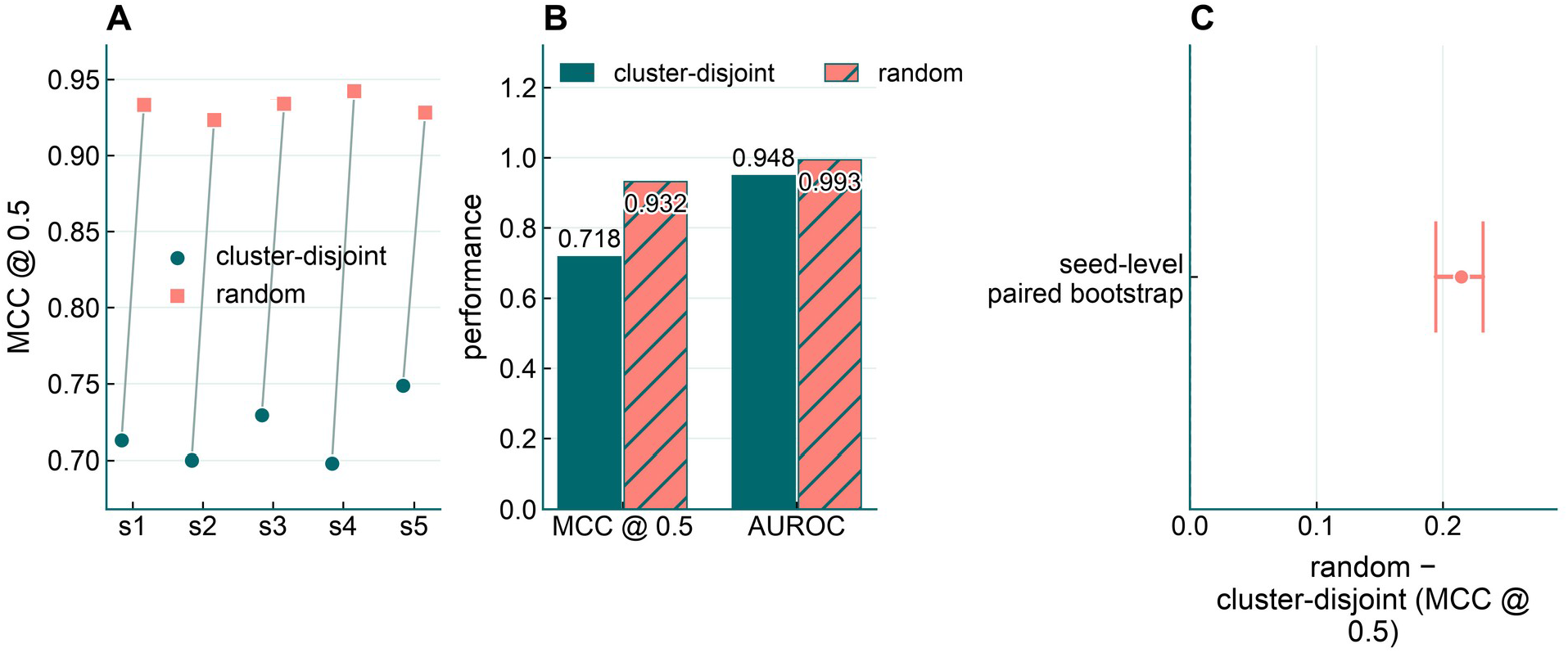
Classification performance under random and cluster-disjoint splits. (A) Test MCC at threshold 0.5 for five matched training seeds on fixed cluster-disjoint and random DRAMP partitions. (B) Mean per-seed MCC and AUROC calculated from mean prediction probabilities across the five runs. (C) Mean paired MCC difference, calculated as random minus cluster-disjoint, with a 95% bootstrap interval across the five matched seeds.

### Split comparisons across additional dataset configurations

The DRAMP cohort contains 26,408 sequences in 9,452 homology clusters. Under a random split, 75.6% of test sequences shared a cluster with a training sequence. We repeated the split comparison on three additional dataset configurations (S1 Table). Exact matches in DRAMP accounted for 79.4% of Veltri positives and 79.1% of AMP Scanner v2 positives. All 2,776 AMP Scanner v2 sequences also occur in the Veltri dataset with the same labels. The two versions assess sensitivity to benchmark composition within a shared sequence collection.

The random-minus-cluster MCC differences were −0.01769 for APD6, −0.04634 for Veltri and +0.05682 for AMP Scanner v2, compared with +0.21441 for DRAMP. Veltri and AMP Scanner v2 had similar train–test cluster overlap under random splitting (13.7% and 12.7%), yet their MCC differences had opposite signs. With the cluster-disjoint split, the APD6 benchmark already reached an AUROC of 0.99848. Its positives and negatives came from different databases, so the classifier could have used features linked to database source. The near-ceiling AUROC left little room to detect an increase under random splitting.

### External evaluation on DBAASP and APD6

On 44,803 DBAASP peptide–organism pairs, the task-3 model’s Spearman *ρ* was 0.367 (Fig 4, Table 4, S10 Fig, S11 Fig). Grouping the pairs by linked article year gave *ρ* = 0.422 through 2014 (17,255 pairs), 0.374 for 2015–2019 (10,380 pairs) and 0.311 from 2020 onward (17,134 pairs). The three groups contained 4,239, 2,713 and 4,353 distinct sequences and covered 197, 161 and 138 organisms, respectively. *Escherichia coli* and *Staphylococcus aureus* were the two most frequent organisms in every group. A separate retrospective analysis of the 2023-and-later subset gave *ρ* = 0.241 (8,628 pairs, 2,222 distinct sequences). The latter period overlaps the 2020-and-later group. Per-organism *ρ* across 21 species ranged from 0.237 to 0.419. Adding DBAASP records to training improved the same temporal test by about +0.078, while global centroid distance to the training set had weak associations with year ( *ρ* =-0.036 and - 0.149 for two centroid variants). Local k-nearest-neighbour distance and ensemble variance both rose with article year, which is consistent with reduced local coverage in the later strata. The temporal analysis is retrospective.

**Table 4.** External evaluation and reliability metrics.

| Evaluation | n | Result |
|---|---|---|
| DBAASP peptide–organism pairs | 44,803 | *ρ*=0.367 |
| Article-year groups | — | *ρ*= 0.422, 0.374 and 0.311. The overlapping 2023-and-later subset gave *ρ*=0.241. |
| Per-organism ρ (21 species) | — | 0.237 to 0.419 |
| APD6 strict positives against DRAMP negatives | 551 | AUROC 0.9833 |
| APD6 against external hard negatives | 1,783 | AUROC 0.9018; specificity 0.6097 |
| Length-matched APD6 arm | 402 | AUROC 0.9047; specificity 0.6443 |
| Expected calibration error | — | 0.0627 |
| Ensemble variance predicting errors | — | AUROC 0.845 |

**Fig 4.**
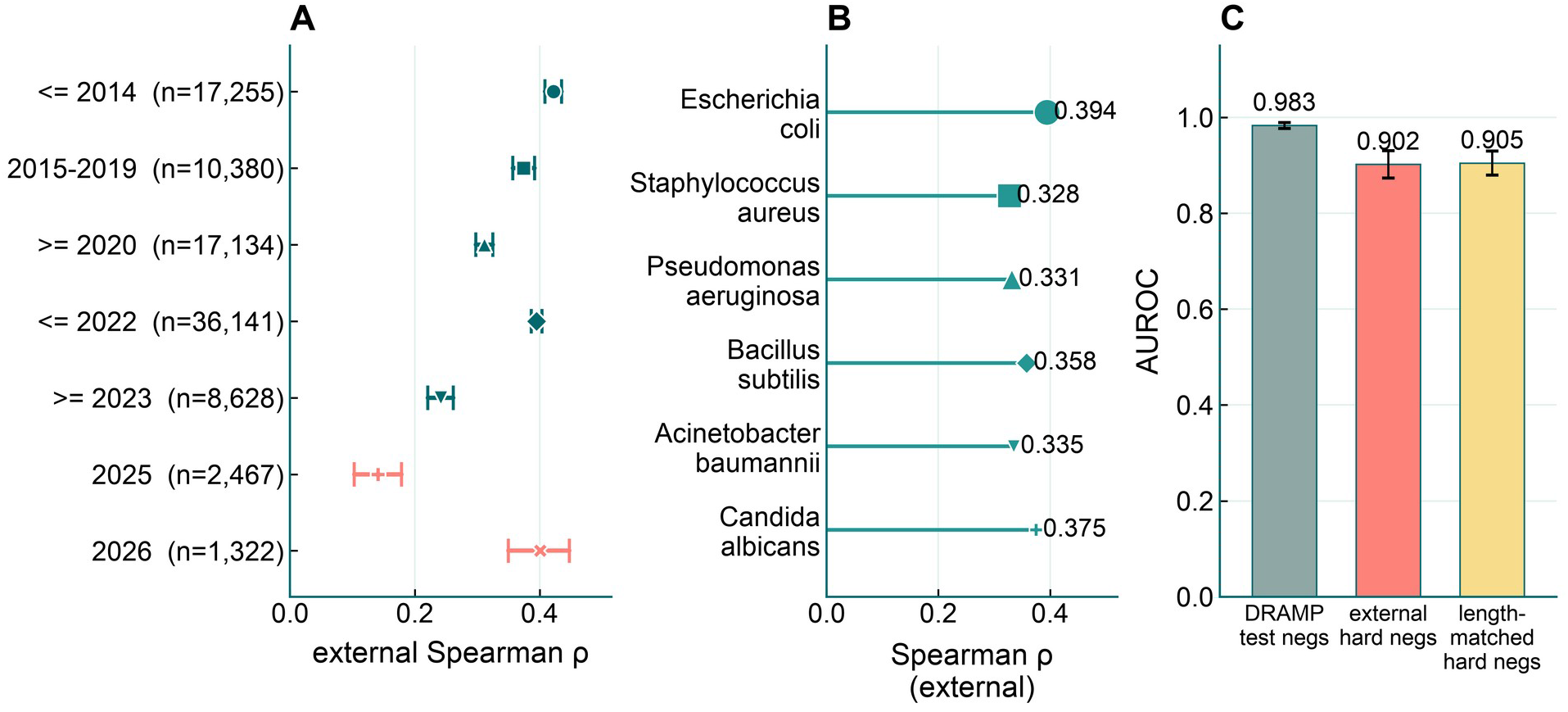
Limits of cross-database generalisation. (A) Spearman correlations between observed and predicted pMIC in DBAASP article-year strata, with 95% bootstrap confidence intervals and peptide– organism pair counts. The 2023-and-later stratum overlaps with the 2020-and-later stratum. (B) External Spearman correlations for the six organisms with the most evaluated pairs. (C) Mean AUROC for APD6 positives evaluated against DRAMP test negatives, external hard negatives or length-matched hard negatives. Error bars show standard deviations across the five model runs.

The AUROC of 0.9833 was measured on APD6 positives paired with DRAMP test negatives. The 44,769-pair complete-case analysis used the years of articles linked to the DBAASP records.

We evaluated the same APD6 [28] strict positives against two negative sets (S4 Fig). AUROC was 0.9833 against the frozen DRAMP test negatives and 0.9018 against 1,783 external hard negatives. Specificity against the external negatives was 0.6097 at the validation threshold. Restricting the comparison to 402 length-matched positives gave AUROC 0.9047 and specificity 0.6443. The external set includes cell-penetrating and haemolytic peptides with possible antimicrobial activity, so these specificity estimates are conditional on the assigned labels.

The AUROC difference was 0.08149 (95% CI 0.07196 to 0.09181, 2,000 paired stratified bootstrap replicates). The AUROC intervals were 0.97692 to 0.98852 with DRAMP negatives and 0.88879 to 0.91402 with external negatives. These intervals reflect resampling of the evaluated peptides with the five fitted models held fixed. Per-seed AUROCs agreed with the original runs to the reported precision.

### Calibration, error detection and model ablations

Calibration on the held-out task-1 test set was close to nominal (expected calibration error 0.0627; calibration slope 1.048) (Fig 5), using binned estimates of calibration error [29]. Distance to the global training centroid, however, failed to identify errors, whereas the variance across the five-seed ensemble did: variance predicted misclassification with AUROC 0.845 and tracked model disagreement at *ρ* = 0.948. Ensemble variance also rose with article year ( *ρ* = 0.433), suggesting that it could flag changes in this cohort without activity labels.

**Fig 5.**
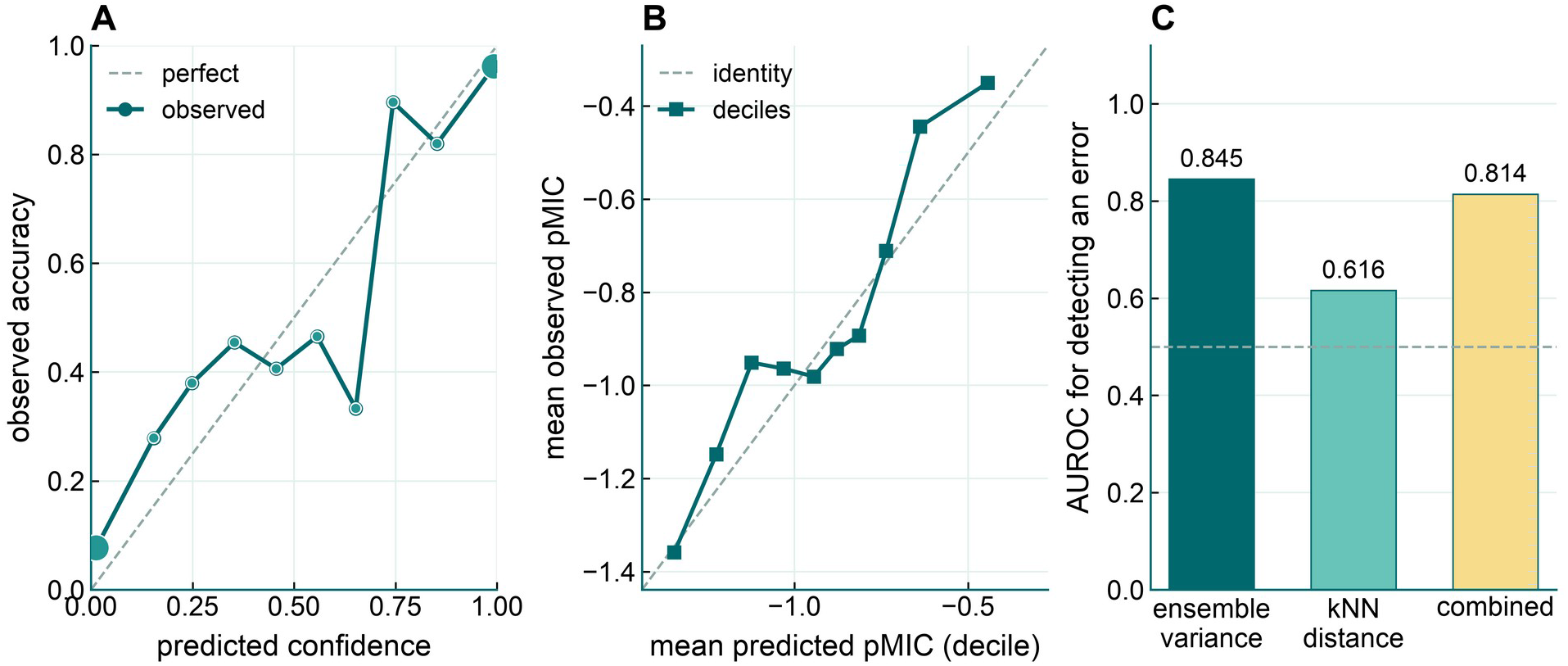
Calibration and reliability diagnostics. (A) Observed accuracy by predicted confidence bin for task 1. Marker area is proportional to the bin count. (B) Mean observed and predicted pMIC by prediction decile for task 3. In A and B, the diagonal marks perfect calibration. (C) AUROC for detecting task 1 classification errors using prediction variance across the five training seeds, distance to the nearest training neighbours, or the combined score. The dashed line marks AUROC 0.5.

The architectural additions tested under the same protocol did not yield a stable benefit (S9 Fig). A molecular-graph branch changed pooled performance by +0.0041 and a chirality channel by +0.0067, both with confidence intervals spanning zero; implementation-to-implementation variation exceeded either effect. Concatenating explicit descriptors reduced performance by 0.027; explicit modification flags changed internal performance by +0.0015 and external performance by −0.028. Multitask training moved task-3 *ρ* from 0.390 to 0.203, and three ablations of the shared architecture reproduced the direction. These ablations estimate the effects within the capacity-controlled architecture tested here.

### Simple baselines and interpretation of peptide features

Two strong simple baselines changed how we read the multi-task results (Fig 6, S5 Fig, S6 Fig, S7 Fig and S8 Fig). For eight-class activity-type prediction, a logistic-regression probe on the same frozen ESM-2 [30] embeddings reached macro-AUROC 0.7842 (95% CI 0.7556 to 0.8096), whereas the shared-trunk MLP head used in the multitask experiment reached 0.648 ± 0.016 with an independent rerun at 0.698 ± 0.022. The shared-trunk estimate comes from the capacity-controlled multitask comparison. For MIC regression, the ESM-2 random forest in Fig 6B achieved a mean test Spearman ρ of 0.38452 across three seeds with organism encoding and 0.36597 without it. Ridge regression using amino-acid composition and organism encoding achieved ρ = 0.3637 (95% CI 0.308 to 0.420). Composition and organism information carry most of the signal. In the scale control, ESM-2 150M scored higher in all five seeds. Mean MCC was 0.75143 ± 0.00970 for the 150M encoder and 0.71778 ± 0.01915 for the 650M encoder, a difference of 0.0337. Data-scale learning curves for the classification and regression tasks are shown in S3 Fig.

**Fig 6.**
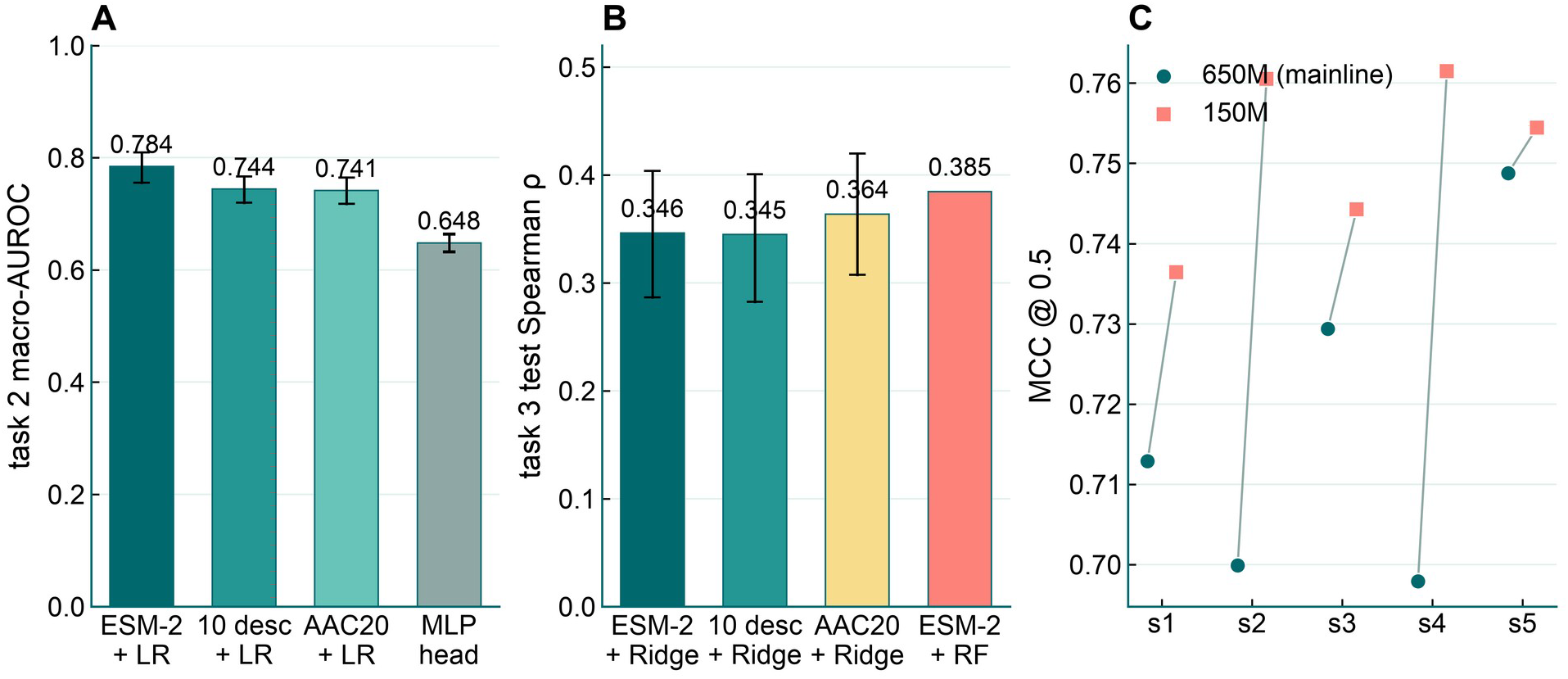
Simple baselines and feature controls. (A) Test macro-AUROC for task 2. The logistic regression probes use ESM-2 embeddings, 10 physicochemical descriptors or 20 amino-acid composition features (AAC20). Capped intervals show 95% bootstrap confidence intervals for the probes and the standard deviation across seeds for the multilayer perceptron (MLP) reference. (B) Test Spearman correlation for task 3. Ridge models use the indicated peptide features and organism encoding. Their error bars show 95% bootstrap confidence intervals. The ESM-2 random forest (RF) reference includes organism encoding and shows the mean across three seeds (0.38452, displayed as 0.385). (C) Paired test MCC at threshold 0.5 for the ESM-2 650M and 150M encoders across five training seeds.

The interpretability analyses are group-level observations (Fig 7, Fig 8, S12 Fig). Among 315 paired DBAASP peptides, C-terminal amidation was associated with a median ΔpMIC of +0.155 ( *p* = 1.7 *×* 10^- 5^); among 169 pairs, N-terminal acetylation was associated with −0.166 ( *p* = 0.0039). Experimental mutational studies have examined the effects of amidation and sequence substitutions on activity and selectivity [31,32]. In our model, alanine-scanning attribution enriched cationic K/R/H hotspots 1.46-fold ( *p* = 4.0 *×* 10^-7^), while agreement between gradient-based attribution and the scan was weak. Haemolysis classification on DBAASP records [20] reached *AUROC* 0.880 *±* 0.011, and the selectivity index correlated with net charge and hydrophobic moment. The corresponding feature tests showed no consistent improvement in prediction.

**Fig 7.**
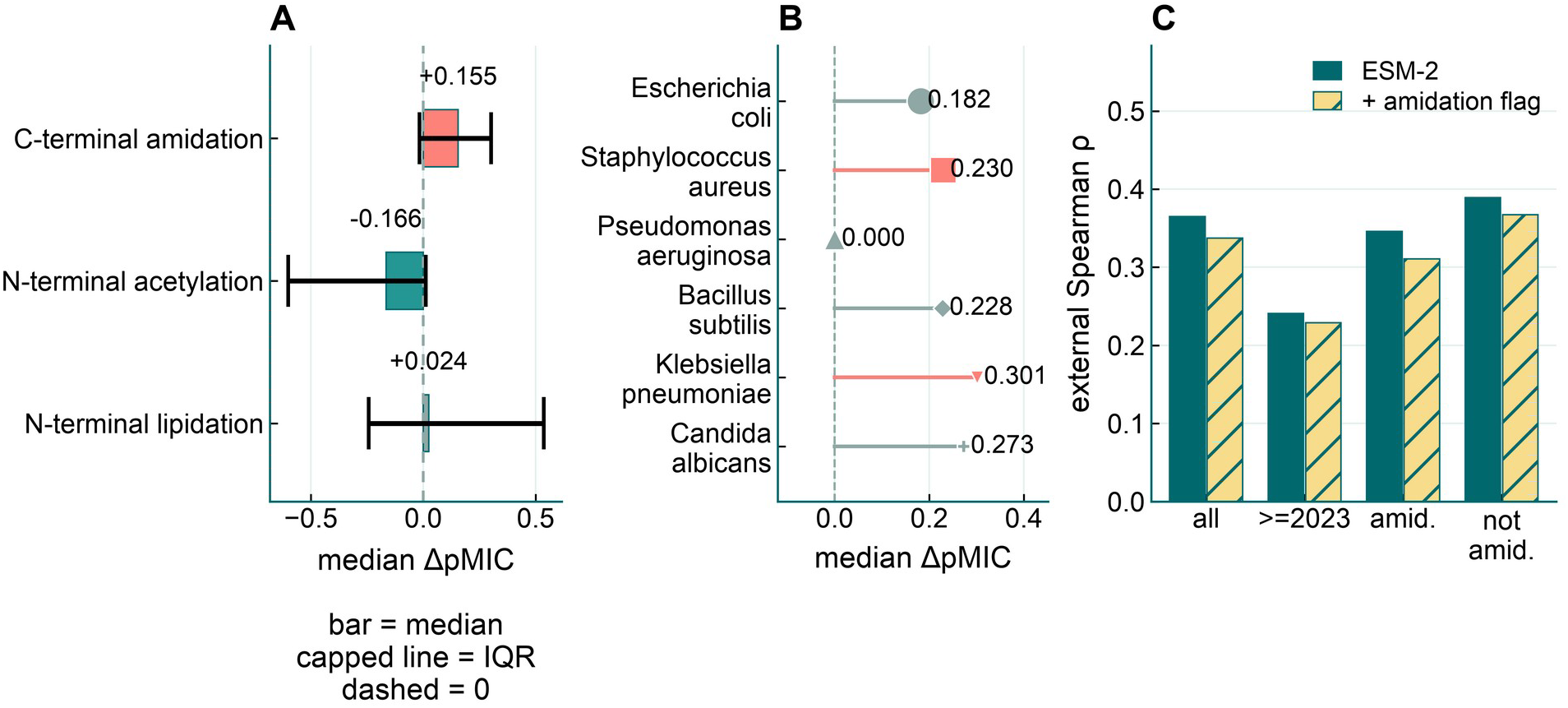
Paired modification effects and prediction checks. (A) Median paired change in pMIC by modification type. Capped lines span the interquartile range, the dashed line marks zero, and coloured bars indicate two-sided sign-test p < 0.05. (B) Median paired change for C-terminal amidation in the six organisms with the most pairs. Marker size reflects the pair count, and coloured markers indicate p < 0.05. (C) External Spearman correlation with and without an explicit amidation indicator added to ESM-2 features, shown for all pairs, pairs from 2023 onward, amidated peptides and non-amidated peptides.

**Fig 8.**
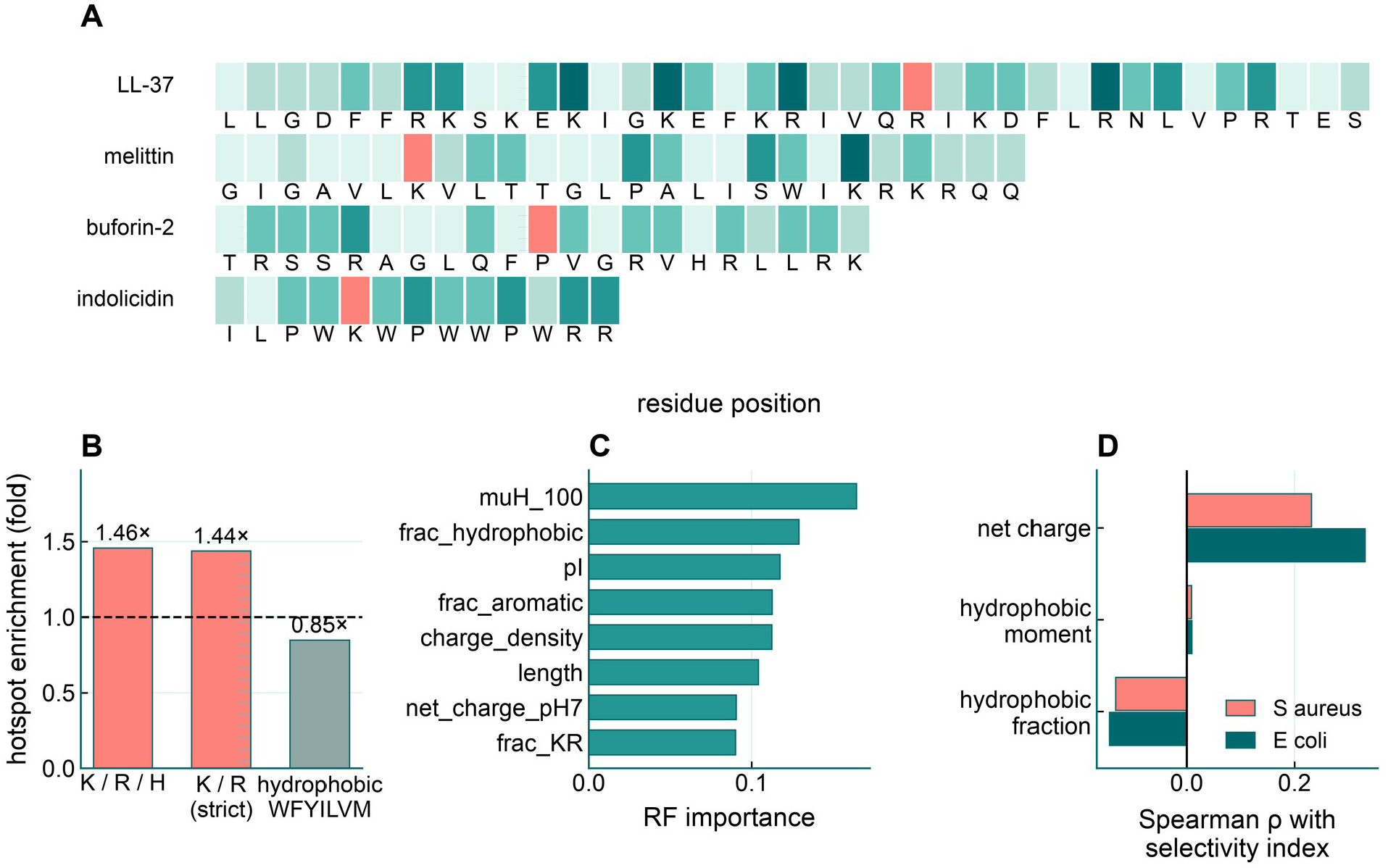
Residue attribution, haemolysis and selectivity. (A) Magnitudes of prediction changes after in silico alanine substitution at each residue of four example peptides. Colour intensity is scaled within each peptide. (B) Enrichment of the indicated residue groups among positions with the largest prediction changes. The dashed line marks no enrichment. (C) Feature importance in the random forest haemolysis classifier. The annotated AUROC is the mean ± standard deviation across five cross-validation folds. (D) Spearman correlations of net charge, hydrophobic moment and hydrophobic fraction with the selectivity index for Staphylococcus aureus and Escherichia coli.

## Discussion

Evaluation design had a substantial effect on AMP performance estimates. On DRAMP, the MCC difference of 0.214 between random and cluster-disjoint splitting exceeded the gains from the architectural additions tested in this study. With 75.6% of randomly assigned test sequences sharing a homology cluster with training sequences, the random partition evaluated predictions in a setting with extensive sequence overlap. The comparison places the architectural results in context. Model improvements need to be assessed against the variation introduced by the choice of training and test sequences.

The APD6 experiment isolates the contribution of test-negative selection more directly. Keeping the fitted classifiers and positive peptides fixed, we observed an AUROC decrease of 0.08149 when DRAMP negatives were replaced with external hard negatives. The paired bootstrap 95% confidence interval for the decrease was 0.07196 to 0.09181. A high score against one negative set therefore provides limited evidence about discrimination against peptides drawn from another source. For AMP screening, the composition of the candidate pool is part of the evaluation question. Database source, peptide length and amino-acid composition can all affect the difficulty of that question [10,13,14].

The additional split comparisons show that the DRAMP effect varies across benchmark configurations. Random-minus-cluster MCC differences ranged from −0.04634 to +0.05682, with opposite signs in the two overlapping Veltri-derived datasets (S1 Table). Their similar random-split cluster-overlap rates, 13.7% and 12.7%, accompanied different performance changes. These results make the composition of each benchmark central to interpretation. Train– test homology, the examples assigned to each partition and the separability of the positive and negative sources jointly determine the observed score. Reporting the clustering rule and measured overlap alongside negative-set provenance allows readers to assess which published evaluations [1,23,24,25,26] address comparable prediction settings.

External MIC evaluation extends this assessment to transfer across databases. Spearman correlation was 0.367 over 44,803 DBAASP peptide–organism pairs, and ranking agreement declined in later article-year strata. Local neighbour distance and ensemble variance increased with article year even though distance to the global training centroid changed little. This pattern is consistent with reduced local coverage of later records. Differences in organism composition, assay conditions and curation practices offer additional explanations [11,21,22]. The data audit also identifies constraints on multimodal modelling. Sequence–structure agreement differed across DRAMP subsets, while repeat MIC measurements had an estimated consistency of approximately 0.82. These properties affect the available inputs and the interpretation of predictive agreement.

The split comparisons use one fixed partition per scheme and dataset. The schemes change both training and test membership. The score differences can therefore reflect both train–test homology and partition difficulty. Variation across five training seeds characterises fitting variability within these partitions. The two Veltri-derived datasets share sequences and represent configurations of a common benchmark. Overlap with UniRef50 pretraining data was not audited directly, and the 150M versus 650M scale comparison changed both encoder size and embedding dimension.

The analyses use existing in vitro activity and MIC records, with most peptide–organism labels derived from a single measurement. The repeat-measurement consistency estimate applies to the subset with repeats. Stereochemical substitutions can affect AMP activity and selectivity [33,34], while the ESM-2 sequence inputs used here do not distinguish D- and L-residues. The DBAASP subset provided no matched D/L pairs among 699 D-amino-acid peptides (S12 Fig). Sparse activity classes and labels derived from page parsing further limit the secondary multi-label analysis. Article-year comparisons are retrospective, including the overlapping 2023-and-later subset and the imprecise rebound in 2026. Prospective evaluation with prespecified MIC extraction and a consistent assay protocol would allow a more direct assessment of temporal transfer.

AMPBenchAudit makes the evaluation choices behind each score inspectable through its audited records, explicit partitions and released analysis code. The controlled negative-set comparison and the variation in split effects identify concrete conditions that AMP benchmarks need to report. This supports model comparisons tied to a defined sequence distribution and screening task.

## Materials and Methods

### Data sources and audit

DRAMP 5.0 FASTA and SMILES records were parsed at record level [17]. Sequence–graph isomorphism between the SMILES-derived structure and the FASTA sequence was evaluated separately for general and stability subsets; the pooled figure was recorded but excluded from modelling decisions because it is dominated by patent-derived records. We used APD6 [28] and DBAASP [20,21] for external evaluation of the main DRAMP-trained models. A separate training experiment added DBAASP records. We parsed DBAASP concentration strings to recover units because the floating-point activity field does not reliably retain them.

### MIC parsing and endpoint definition

MIC values were extracted from the free-text Target_Organism field, restricted to MIC endpoints, converted to micromolar units and aggregated across repeated measurements under four rules; agreement between rules was *ρ≥* 0.978. The regression target was *p_mic_* =- log_10_ *MIC* in *μM*. IC50, MBC, EC50, IC90, LD50 and MFC endpoints were counted in the audit and excluded from the regression cohort. Spearman–Brown corrected consistency was estimated from records with two or more measurements.

### Homology-aware partitioning and negatives

Sequences were clustered by single linkage at a 0.80 sequence-identity threshold calculated from global alignments [27], and clusters were assigned as whole units to train, validation or test so that no cluster spans two partitions. The negative set contained 13,204 rows representing 5,886 unique sequences. We checked length and amino-acid composition for potential shortcuts. Task-1 splits contain 26,408 sequences (test 2,549). Task-3 splits contain 2,378 sequences (test 238). The two split files are not interchangeable.

### Tasks, model and training

Three tasks were defined: binary AMP/non-AMP classification, eight-class activity-type prediction with a masked multi-label formulation (6,660 masks, no mask violations), and pMIC regression. The main representation was frozen ESM-2 650M embeddings (esm2_t33_650M_UR50D) [30] with a shallow prediction head; the same representation was reused for all downstream analyses. All main experiments used five random seeds, and the primary task-1 metric was MCC at a fixed threshold of 0.5. Validation-tuned thresholds are reported in the supplement but are not the primary operating point, because the four public tools have widely separated optimal thresholds (0.15, 0.675 and 0.95 among them). We used the 650M encoder across the downstream analyses to keep the representation consistent. The scale control compared it with the 150M encoder. For the architecture ablations, we computed paired differences between runs with the same training seed. Four of the eight activity labels were derived by page-parsing rules that may undercount relative to the official annotation. The rare classes contained 174 antiprotozoal and 33 antiparasitic records and were not learnable in these experiments. Task-2 reruns fluctuated by about 0.05. An early Gate-2 implementation showed nondeterministic scatter-add behaviour, which was resolved by cross-implementation pooling. Task-2 uncertainty is reported using five-seed summaries and pooled cross-implementation intervals. Gate-2 seed-level paired intervals and an internal task-3 sample-level bootstrap interval were not calculated. Task-3 uncertainty is described using five-seed summaries and external and temporal intervals.

### Baseline re-implementation

We ran AMPlify [23], Macrel [24], AMP Scanner v2 [25] and AMP-BERT [26] on the task-1 test partition at a fixed threshold of 0.5. AMPlify had scores in the descriptive evaluation for the full set, while its paired comparison used the rows that joined by DRAMP ID in the paired-analysis input. AMP Scanner v2 used the rows with matched scores in that input. Macrel normalises some sequences, so we report descriptive scores for both sequence-key and ID-key alignment and use the ID-key match for the paired comparison. AMP-BERT matched the full test set. The training-set overlap of the released tools with this test partition was not established. The paired point estimates use mean MCC across the five ESM-2 training seeds. We used a paired bootstrap over matched test sequences to estimate confidence intervals for MCC differences and DeLong’s test [35] to compare AUROCs. Both analyses used the mean predicted probability across seeds for each test sequence. The attempted iAMP-CA2L reproduction did not produce a benchmark score.

### External, temporal and reliability evaluation

The overall DBAASP evaluation used 44,803 peptide–organism pairs. The complete-case bootstrap used 44,769 pairs with observed pMIC, predicted pMIC and article year. The dataset builder recorded the earliest parseable year among the articles linked to each DBAASP entry. Bootstrap replicates sampled peptide–organism rows with replacement. Multiple rows can share a peptide, so the row-level intervals do not account for within-peptide dependence. APD6 strict positives were evaluated against both the frozen DRAMP test negatives and 1,783 external hard negatives, with a length-matched arm where matching was possible. The external set contained 1,238 UniProt signal peptides, 288 natural and 4 non-natural CPPsite peptides, and 253 Hemolytik peptides [36]. We screened these candidates for exact overlap with DRAMP and shared 8-mers with APD6. We assessed calibration using binned expected calibration error [29] and calibration slope. We also examined global centroid distance and local k-nearest-neighbour distance and evaluated variance across the five training seeds as an uncertainty score [37]. Modification effects were estimated from paired DBAASP peptides, attribution from alanine scanning, and safety read-outs from the haemolysis and selectivity-index subsets (S12 Fig). The prespecified prospective MIC extraction remains incomplete, so article-year strata are interpreted retrospectively.

### Replication cohorts and paired bootstrap

We repeated the split comparison using APD6 strict positives paired with external hard negatives and two versions of the Veltri benchmark [25,26]. The larger version came from the AMP-BERT distribution [26]. For the smaller version, we pooled the original training and test files distributed with AMP Scanner v2 [25]. We used the same single-linkage rule, identity threshold and split seeds as for DRAMP. For each dataset, we generated one cluster-disjoint partition and one random partition with matching class counts in the training, validation and test sets. We held each partition fixed across the five training seeds. For the random split, we counted the fraction of test sequences that shared a cluster with at least one training sequence.

For the APD6 negative-set comparison, we sampled peptides with replacement separately from the APD6 positives, DRAMP test negatives and external hard negatives. In each replicate, we used the same positive resample for both AUROCs and applied the same resampling indices to the predictions from all five models. We calculated both AUROCs for each model and averaged the paired differences across models. S1 File provides the replicate count, bootstrap seed and sample-level probabilities.

### Declaration on usage of Large Language Models (LLMs)

During the preparation of this work the authors used ChatGPT 6.1 in order to help with the coding process and to improve language and readability. After using this tool, the authors reviewed and edited the content as needed to ensure the content is accurate and valid. The authors ensure all statements in the article reporting hypotheses, interpretations, results, conclusions, limitations, and implications of the study represent the authors’ own ideas and take full responsibility for the content of the article.

## Data and Code Availability

The raw DRAMP 5.0, DBAASP and UniProtKB inputs included in the AMPBenchAudit archive are available from [Zenodo](https://zenodo.org/records/24268404). The code for data processing, model training and evaluation is available from [GitHub](https://github.com/xiaoniao007-debug/amp-bench-audit). Split assignments and result summaries for the supplementary replication analysis, together with the APD6 negative-set bootstrap results and prediction probabilities, are provided in S1 File.

## Author contributions

**Conceptualization:** Chen Chen and Yanshan Tian.

**Formal analysis:** Hongbo Zhou, Qifu Li and Zishang Gao.

**Software:** Chen Chen.

**Validation:** Hongbo Zhou, Qifu Li and Zishang Gao.

**Writing – original draft:** Chen Chen.

## Supporting information

**S1 Table. Sequence redundancy and performance under random and cluster-disjoint splits.** Clusters were formed by single linkage at a common identity threshold. ΔMCC is the difference between MCC at a threshold of 0.5 under random and cluster-disjoint splitting, averaged over five training seeds. Each set of five runs used a fixed partition.

**S1 Fig. Data-audit detail.** Stratified graph-isomorphism rates show that the pooled value is driven by patent-derived records; mismatch classes and source subsets are drawn separately.

**S2 Fig. Additional statistics for the public predictor comparison.** An independent CPU reproduction, AUROC on matched subsets and tuned-threshold sensitivity complement the fixed-threshold comparison in the main text.

**S3 Fig. Data-scale learning curves.** The classification task saturates early, whereas the regression task keeps improving with additional data.

**S4 Fig. APD6 hard-negative diagnostic.** Length matching does not explain the drop in specificity; the external negatives include cell-penetrating and haemolytic peptides, so the specificity estimate depends on the assigned negative labels.

**S5 Fig. ESM-2 scale control.** Only the backbone and its embedding dimension change between arms. The pretraining-overlap audit was not feasible, so the comparison stays directional.

**S6 Fig. Task 2 per-class behaviour.** All activity classes are shown with their support; measured performance varies with class support.

**S7 Fig. Task 3 feature ablation.** Adding explicit descriptors reduces ranking agreement relative to the frozen representation alone, while a descriptor-only arm shows physicochemistry dominating.

**S8 Fig. Organism-conditioning ablation.** Removing organism one-hot encoding reverses direction for one organism, so the conditioning claim stays modest; paired seed-level intervals are directional only.

**S9 Fig. Multitask and structure ablations.** Multitask training degrades both the regression and classification tasks, and the confidence interval for the structure branch spans zero.

**S10 Fig. Temporal decay and drift diagnostics.** Ranking agreement declines across yearly bins, the later stratum is examined on its own, embedding drift stays flat and model disagreement rises slowly.

**S11 Fig. Per-organism asymmetry.** Data volume correlates only weakly with ranking agreement, calibration error is roughly uniform, and all evaluated organisms are shown.

**S12 Fig. Attribution agreement and D-amino-acid limits.** Gradient-based attribution agrees only weakly with the perturbation scan; no matched D/L pairs were available, so stereochemistry cannot be assessed.

S1 File. Replication results and APD6 negative-set bootstrap. ZIP archive containing split assignments, analysis summaries, bootstrap inputs and results, and SHA-256 checksums.

